# Cement lines are stiffer and harder than bone but exhibit different mineral–mechanics relationships due to thicker and shorter mineral particles

**DOI:** 10.1101/2025.08.25.672154

**Authors:** Astrid Cantamessa, Victoria Schemenz, Stéphane Blouin, Timothy Volders, Shahrouz Amini, Maximilian Rummler, Manfred Burghammer, Jiliang Liu, Andrea Berzlanovich, Richard Weinkamer, Wolfgang Wagermaier, Markus A Hartmann, Davide Ruffoni

## Abstract

The remarkable mechanical performance of bone arises from its complex hierarchical structure and from the presence of numerous internal interfaces joining the different components. Formed during bone remodeling, the cement line (CL) is a thin interface surrounding osteons. Although often neglected due to its small size, the CL has been suggested to play an important role in bone fracture toughness. However, there is an ongoing debate about its specific structure and mechanical behavior. Here, we investigate the composition-mechanics relationship at the CL and surrounding bone in human osteons using multiple methods in a correlative manner. We found that the CL is stiffer and harder than adjacent osteonal tissue. However, the CL requires more mineral than bone to attain the same stiffness and hardness. Analyzing the nanoscale properties of the mineral, we found thicker but shorter particles at the CL. Using a mechanical model, we interpreted the lower aspect ratio of the mineral particles as a less effective way to reinforce the collagen matrix. Casting our findings into a computational model, we questioned the possible protective role of the CL, which is not a soft interface as traditionally believed.

## Introduction

Bone is a load bearing material which needs to be stiff and tough.^1^ The stiffness is required to prevent excessive deformation under daily loadings, which can even be higher than body weight.^2^ The toughness is needed to prevent low-energy impacts from causing bone fractures. This is particularly important because bone must be also relatively lightweight and is, therefore, not overdesigned: the margin between physiological and failure loads is not as high as in most engineering structures.^3, 4^ While healthy bone effectively combines those conflicting requirements, aged and diseased bone becomes weaker and more susceptible to fractures.^1, 5, 6^

It is fascinating that the remarkable mechanical behavior of bone is obtained using limited constituents: bone is essentially composed of a mineral phase (stiff but brittle), a collagen-based matrix (tough but compliant) and water.^3^ The constituents are organized into a complex three-dimensional hierarchical structure, which offers the opportunity to adapt each level of the structure to different requirements.^7^ A hierarchical structure also implies that building blocks of different dimensions have to be assembled, generating numerous internal interfaces at many length scales.^8^ Therefore, bone mechanical behavior, especially fracture resistance, depends not only on the building blocks but also on the properties of the interfaces joining them.^5^

At the nanoscale, tiny mineral particles reinforce the collagen molecules: ionic bonds form at the interface between these elements, linking amino acids from the protein to calcium ions from the mineral.^9^ Several collagen molecules assemble into mineralized collagen fibrils and the interfacial space between fibrils is occupied by non-collagenous proteins and proteoglycans. These interfaces are critical for the deformation of bone at the nanoscale: load sharing between mineral and collagen occurs through interface shear and sliding,^10, 11^ causing the breaking of sacrificial bonds to dissipate energy and increase bone toughness.^12^ At the next hierarchical level, mineralized collagen fibrils are arranged into fiber bundles, which appear in different structural motifs, including the more disorganized woven bone and the ordered lamellar bone.^13^ A distinct feature of lamellar bone is the organization in sheets of parallel collagen fibers, which are stacked under different orientations to form the so-called rotated plywood structure.^14, 15^ Numerous works have shown that lamellar bone can enhance fracture toughness by several mechanisms, including a periodic modulation of elastic properties^16^ and the deviation of cracks at the interface between lamellae.^17^ Lamellar bone is further organized in osteons, which are characteristic structures of cortical bone, where a central Haversian canal (hosting blood vessels) is surrounded by a concentric arrangement of lamellae.^18^ Secondary osteons, formed during bone remodeling, are bordered by a thin interface known as the cement line (CL). Although discovered more than 150 years ago, the CL is still under-investigated in comparison to lamellar bone,^19^ with open questions on CL composition, mechanical behavior and function. Owing to their tiny dimensions (only 1-2 µm in width in longitudinal cross sections), CLs are difficult to be characterized quantitatively; it is also challenging to distinguish their properties from those of surrounding bone.

The present work focuses on the CL, which is the first material layer of a new osteon deposited during a remodeling event.^20–22^ We know that the CL is enriched with osteopontin (a non-collagenous protein),^23^ that it is often more mineralized and follows a different mineralization process than surrounding bone.^24–26^ There is some limited evidence that the CL has lower collagen content^24^ and different mineral characteristics^27^ than osteonal bone. Considering its biological function, the CL may act as a rigid substrate that facilitates the initial assembly and alignment of many osteoblasts, which is required for the formation of organized lamellar bone.^28^ Mechanically, the CL has been suggested to influence crack propagation, contributing to bone toughness. This putative role comes from the observation of crack deflection at the outer boundary of osteons, a mechanism which strongly depends on crack length and trajectory.^29–31^ Additionally, (micro)cracks are more frequently detected in the outer interstitial bone than within the osteons.^32, 33^ Understanding the possible mechanical contribution of CL in bone fracture resistance requires knowledge of CL material properties: in analogy with fiber reinforced composites,^34^ a soft CL may protect the osteon by arresting cracks, whereas a stiff CL may deflect the crack around the osteon^5^. Several computational works have investigated the interplay between microcracking and CL in great details,^35–38^ yet making assumptions on CL properties given the lack of experimental data. Only a very limited number of experimental studies have focused on CL mechanical characteristics, suggesting a softer response compared to the surrounding bone.^39, 40^ However, this appears hard to reconcile with the hypermineralization of the CL, given the close relationship between mechanical properties and mineral content.^11, 41, 42^

In this work we aim: i) to quantify the local mechanical properties of the CL and the composition-mechanics relationships at the CL and surrounding bone, and ii) to measure the nanoscopic mineral characteristics of the CL for elucidating its mechanical performance. By employing several techniques with sub-micrometer resolution in a spatially resolved and correlative manner we wish to advance our knowledge of this enigmatic interface.

## Results and Discussion

Using a multimodal approach, we obtained quantitative two-dimensional maps of mechanical properties, composition and mineral characteristics of the CL and surrounding bone in adult human osteons (an overview of the analyzed samples and methods is given in Supplementary Information, Table S1). We first present findings on mechanical and compositional aspects obtained combining nanoindentation (nIND), quantitative backscattered electron imaging (qBEI) and high-resolution scanning electron microscopy at the same locations (Fig. 1A-E). We complement this analysis by characterizing mineral particles using synchrotron micro-focus scanning X-ray scattering (Fig. 1F-H). Micromechanical modeling is subsequently employed to link the mineral features to the mechanical behavior of the CL. Finally, a proof-of-concept damage-based finite element analysis is proposed to question the mechanical role of the CL.

**Fig. 1:**
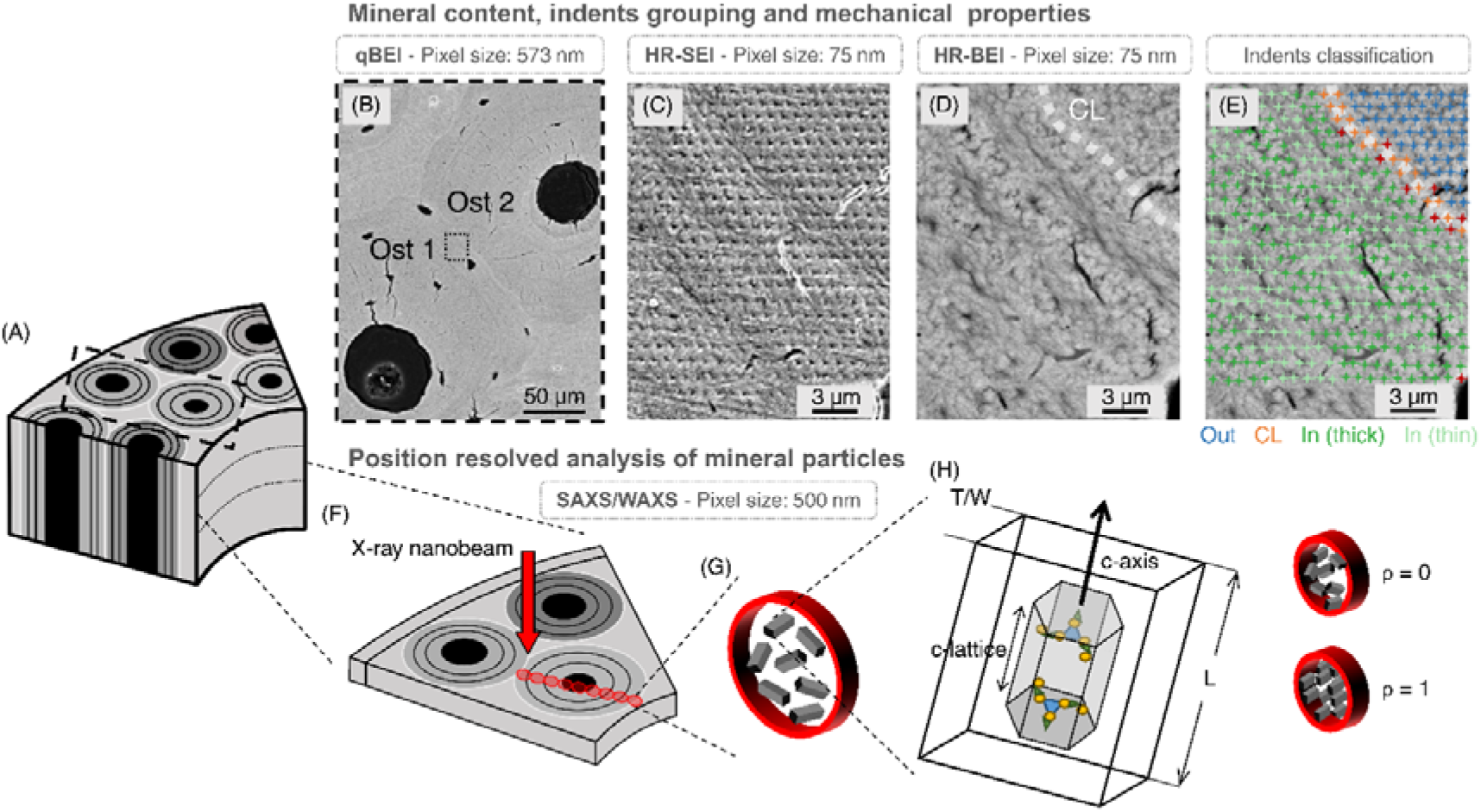
Multimodal analysis of the CL and surrounding bone. (A) Schematic representation of cortical bone. (B) Mineral content map (obtained with quantitative backscattered electron imaging, qBEI) showing two osteons (Ost 1 and Ost 2), the CL and interstitial bone. (C) Magnified view (based on high resolution secondary electron imaging, HR-SEI) of the indented region across the two osteons (inset in B), visualizing indent imprints. (D) High resolution backscattered electron imaging (HR-BEI) of the same region shown in (C) highlighting the CL (dotted line) and the bone lamellae which are recognizable by different textures. (E) Indents are classified as placed in the CL (orange), in the corresponding osteons (In, dark green or bright green depending on the lamella type), or in the surrounding outer bone (Out, blue). Indents located within cracks/pores and not directly falling in the CL were excluded from the analysis (red). (F) Extraction of a thin slice to perform position-resolved analysis of mineral particles using small- and wide-angle X-ray scattering (SAXS/WAXS). Schematic representation of (G) bone mineral particles and (H) different mineral parameters including particle thickness (T or W), particle length (L), crystal lattice constant (c-lattice) and degree of orientation (ρ).

### The cement line is stiffer and harder than the corresponding osteon

Figure 2 shows typical high-resolution maps of local mechanical properties, mineral content, and collagen orientation obtained across the border between two osteons. Qualitative observations reveal a spatial modulation of indentation modulus (Fig. 2B) and hardness (Fig. 2C) in the most recently deposited osteon (Ost 1), as well as higher mechanical properties in the CL and in the older osteon (Ost 2). Within Ost 1, variations in mechanical properties mirror the lamellar organization inferred by SHG imaging (Fig. 2E): bright lamellae in SHG, indicative of (predominantly) in-plane collagen fibers, correspond to a more compliant and softer mechanical behavior; dark lamellae in SHG, having mainly out-of-plane fibers, correspond to stiffer and harder locations. Variations in the local mechanical properties may also depend on the presence of ordered and disordered material in lamellar bone.^43, 44^ The region-matched map of the mineral content (Fig. 2D) also shows variations at the lamellar level, albeit much less evident. A clear feature of Fig. 2D is the increase in mineral content at the CL, which is consistent with higher indentation modulus and hardness compared to the corresponding osteon. Elevated mechanical properties are maintained in the outer region belonging to the neighboring osteon (Ost 2), although the mean mineral content is lower than in the CL and rather similar to Ost 1 (Fig. 2D). A possible explanation of this trend could be the predominance of out-of-plane fibers, presumed by the missing SHG signal (see also Supplementary Information, Fig. S4), which lead to a stiffer behavior.^45, 46^

**Fig. 2:**
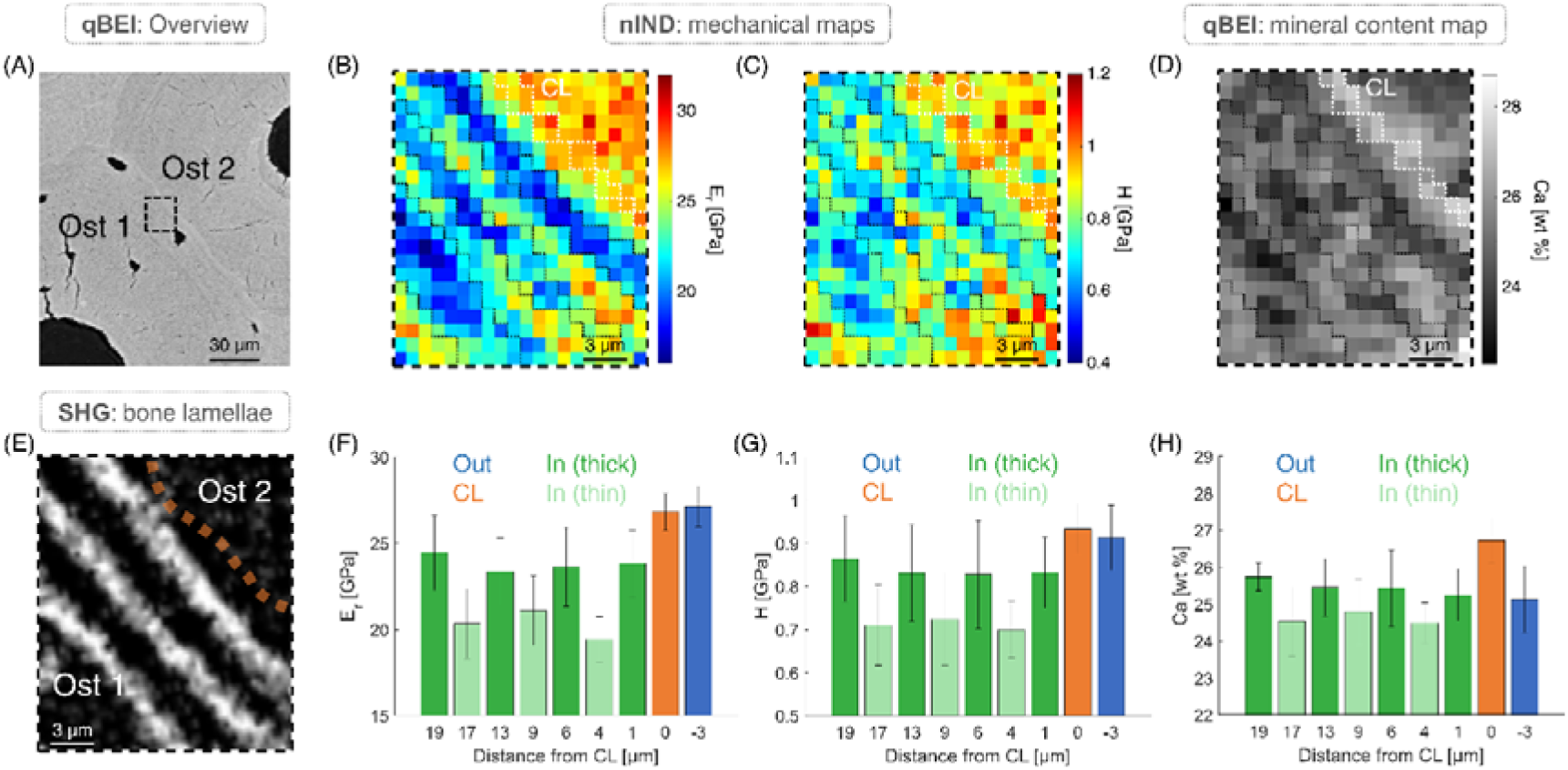
Correlative analysis of mineral content and mechanical properties. (A) Overview qBEI showing two osteons (Ost 1 and Ost 2) having a similar mineral content and separated by a brighter CL. The indented area is highlighted by the black dashed frame. Two-dimensional maps of (B) indentation modulus Er, (C) hardness H, and (D) calcium content Ca. The dotted lines contour the position of the CL (in white) and of the bone lamellae (in black). (E) SHG image of the same region highlighting the lamellar structure of Ost 1 and the location of the CL (dashed line). (F-H) Bar plots showing mean and standard deviation of Er, H, and Ca in the CL (orange), within the bright and dark lamellae of the corresponding inner osteon (In, light and dark green) and in the external outer bone (Out, blue).

The precise localization of the indents in the mineral content maps (see Methods) allowed us to calculate mean stiffness (indentation modulus E_r_, Fig. 2F), hardness (H, Fig. 2G) and calcium content (Ca, Fig. 2H) in different regions. The CL is the stiffest (E_r_ = 26.82 ± 1.08 GPa) and the hardest (H = 0.93 ± 0.06 GPa) feature of Ost 1, being about 20% stiffer (p < 0.01) and 18% harder (p < 0.01) than the corresponding osteon. A pronounced mechanical contrast is observed between bright (E_r_ = 23.66 ± 2.12 GPa, H= 0.83 ± 0.11) and dark (E_r_ = 20.27 ± 1.92 GPa, H= 0.71 ± 0.09) lamellae within Ost 1, corresponding to a 15.4% difference in stiffness (p < 0.01) and 15.6% in hardness (p < 0.01). The mineral content also shows variations between dark (Ca_Mean_ = 25.44 ± 0.86 wt %) and bright (Ca_Mean_ = 24.6 ± 0.82 wt %) lamellae, with the difference being around 3% (p < 0.01), and possibly due to orientation effects in qBEI (see Limitations). In line with greater mechanical properties, a pronounced peak in mineral content is observed at the CL (Ca_Mean_ = 26.72 ± 0.62 wt %), exceeding the mean mineral content of the corresponding osteon and of the outer bone by more than 6% (p < 0.01). However, no significant differences were found between the mechanical properties of the CL and the outer bone (p = 0.99 and p = 0.87 for Er and H, respectively). These trends are consistently observed in all samples (Supplementary Information, Fig. S5), with the CL exhibiting always higher mineral content (+8.93%) and mechanical properties (+21.92% for E_r_ and +25.87% for H) compared to its osteon (p < 0.01). Although the CL remains more mineralized than the surrounding external bone (+3.44%, p < 0.01), its stiffness and hardness are comparable to those of the outer regions (0.53% in E_r_, p = 0.9; 3.25% in H, p < 0.01).

The mechanical behavior of the CL uncovered here challenges current knowledge: we found that the CL is stiffer and harder than the corresponding osteon and, at the same time, the CL seems to be mechanically comparable to (older) surrounding bone. In contrast, previous works assessing CL mechanical properties with nanoscale methods, reported that the CL is more compliant and softer than neighboring tissue.^39, 40, 47^ This discrepancy could be due to different samples (human femurs analyzed here versus bovine and ovine bones considered in the literature), testing protocols and conditions. Moreover, owing to the small dimensions, the correct identification of the CL is particularly challenging. A distinctive feature of our approach is that the mechanical characterization of the CL is supported by correlative local measurements of the mineral content as well as by a careful localization of the indents falling in the CL with SEM (Fig. 1). The periodic variations in mechanical properties at the lamellar level seen in Ost 1 align with previous studies^48–53^ and, most likely, reflect different arrangements of the mineralized collagen fibers in dark and bright lamellae.

### The cement line needs more mineral than lamellar bone to attain the same mechanical properties

We further explore the relationship between mechanical properties and mineral content of the CL and neighboring bone by collecting and plotting values of indentation modulus and hardness versus calcium content measured at the very same locations. A correlation is observed between mineral content and mechanical properties (Fig. 3A and Supplementary Information, Fig. S6A), but data appears quite scattered, and the corresponding Spearman correlation coefficients range from weak to moderate. This is not surprising considering the underlying heterogenous arrangement of mineralized collagen fibrils.^54, 55^ Qualitative similarly scattered data were obtained at the bone-tendon^42^ and bone-cartilage^41^ interface. More important, data points corresponding to the inner osteon and the surrounding bone align along a common line (connecting the two means), whereas those from the CL are distinctly shifted toward higher mineral content for a given stiffness. To validate this observation, we binned the indentation modulus (and hardness) and computed the corresponding average calcium content needed to reach the specific mechanical property in each bin (Fig. 3B).

**Fig. 3:**
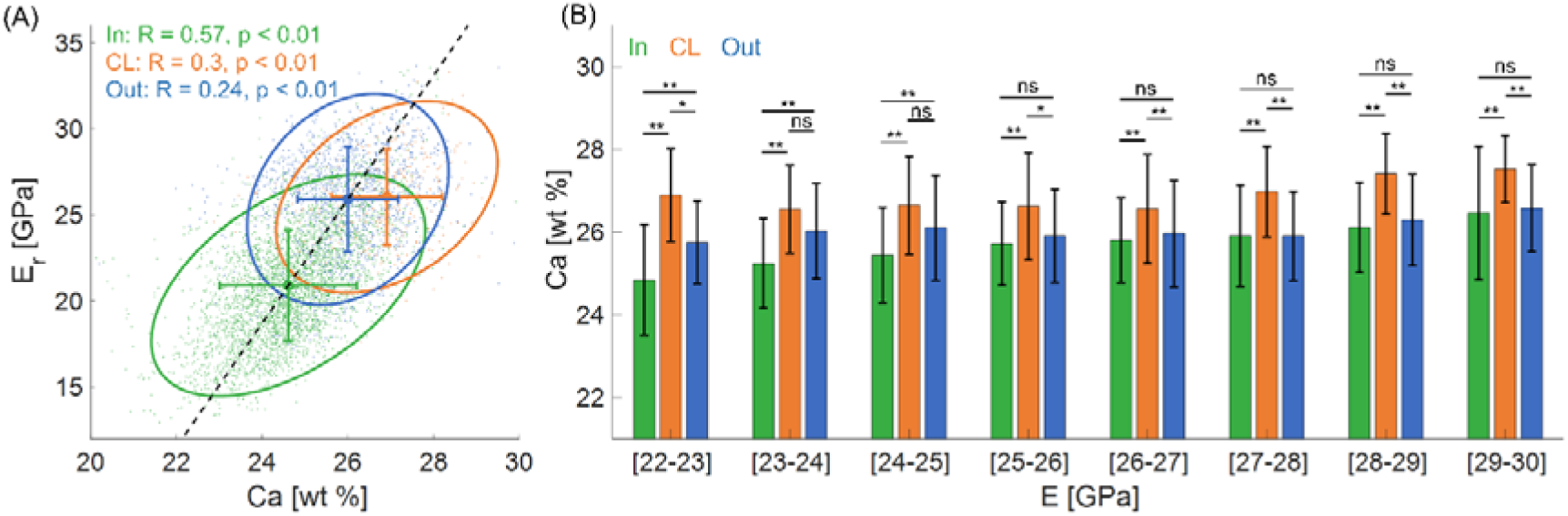
Quantitative relationship between indentation modulus (E_r_) and calcium content (Ca) at the CL, corresponding inner osteon (In), and outer bone (Out). (A) E_r_ versus Ca plot for all samples (total of 4966 indents), the individual Spearman correlation coefficients and p-values are reported. For each group, the scattered data are enveloped by an ellipse representing the 95% confidence interval and centered around the mean. Group means and standard deviations are also displayed. (B) Average mineral content required to achieve the specific stiffness value of each bin, ranging from 22 to 30 GPa (bin width 1 GPa). Error bars represent standard deviations. Statistically significant differences between the data are indicated as follows: *p < 0.05, **p < 0.01, and ns (not significant).

This analysis confirms that, for equivalent mechanical properties, the CL consistently exhibits a higher degree of mineralization. A similar pattern is observed for hardness (Supplementary Information, Fig. S6B). This is a central finding on the mineral content-mechanical property relationship of the CL, which requires a higher calcium content to attain the same stiffness and hardness of the surrounding lamellar bone. This could be due to differences in the collagen-based matrix, mineral crystals or mineral-collagen packing. Considering the mineral phase, there are some indications, albeit limited to a small portion of a single osteon, that the CL may have different mineral characteristics than adjacent bone.^27^ Therefore, we further investigate the property of the mineral crystals to clarify the (nano)structure-mechanics interplay.

### The cement line has thicker and shorter mineral particles than the adjacent bone

High resolution small- and wide-angle X-ray scattering (SAXS and WAXS) are used to analyze the nanostructural properties of the mineral particles, including particle thickness (T and W), length (L), degree of orientation (ρ) and crystal lattice spacing (c-lattice), in a position-resolved manner.

Figure 4 presents two-dimensional maps and line profiles of mineral particle properties across the CL separating two osteons. From the colored maps of zinc intensity (Fig. 4B), we could identify a brighter layer that matched the CL location given by qBEI (Fig. 4A). Zinc is a well-known trace element of bone, and there is evidence that it accumulates at the mineralization front^56^ and at the CL^57^, yet its specific role and possible impact on bone mineralization are still debated. We used the position of the zinc intensity peak to locate the CL on the other images of mineral parameters. Considering the thickness of the mineral particles, the so-called T parameter shows a decrease that colocalizes with the CL (Fig. 4C). As the mineral content of the CL is significantly higher than in the surrounding bone^26^ (Fig. 2H and Supplementary Information, Fig. S7A), we also computed the W parameter, which takes into consideration differences in mineral volume fraction.^58^ As the mineral content was measured with qBEI on a different section than was analyzed with X-ray scattering (about 15-20 µm away), we prefer to correct for mineral content using values averaged over small sectors of these three regions. Therefore, 3 averaged values of mineral volume fraction representative for (i) the CL, (ii) the osteon, and (iii) the older outer bone were used. The W parameter clearly reveals that the CL has thicker mineral particles than neighboring tissue. Similar results are obtained also when using a profile plot of the mineral volume fraction rather than 3 discrete values: the W parameter is higher at the CL while showing a more gradual but noisier pattern (Supplementary Information, Fig. S7). The profile plots of the other parameters (Figs. 4E-G) do not highlight specificities at the CL, one possible reason being a high heterogeneity also evident in the 2D maps (Fig. 4A). One additional feature observable in the spatial maps of mineral characteristics is a local modulation corresponding to the lamellar organization. This underlines that the orientation of the mineralized collagen fibrils influences the observed mineral properties: T and W are mostly orientation independent, while L and ρ depend, to some extent, on orientation (see Limitations).^14, 59–61^ Qualitative similar results are obtained when inspecting other osteons and sections (Supplementary Information, Fig. S8-10).

**Fig. 4:**
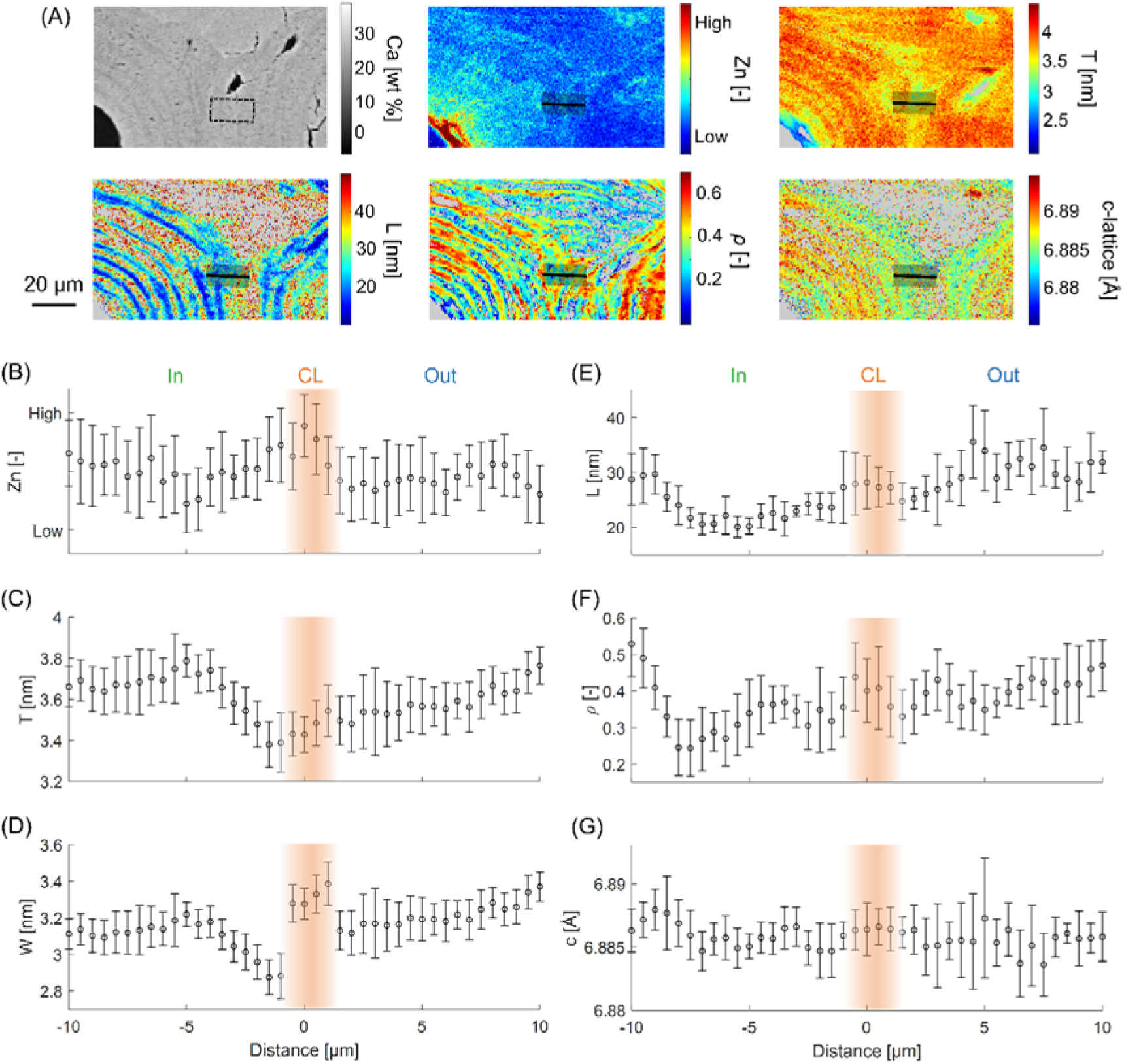
(A) Two-dimensional high-resolution maps of mineral particle characteristics across the CL in Ost 5. Highlighted in the maps (thick black line) is the location across the CL where data is extracted. Profile plots of (B) zinc content and (C-G) particle properties plotted as a function of the distance from the CL. The position and the thickness of the CL are represented by the colored area. Negative distance corresponds to regions inside the osteon and positive values outside. To reduce noise, at each position data are averaged over 20 pixels (shaded areas in A) and displayed as mean values ± standard deviations.

To overcome the challenge of high tissue heterogeneity combined with a limited thickness of the CL, we further compare CL against adjacent bone using frequency distributions based on all collected data (about 14400 pixels, Fig. 5). This comparison corroborates the message that average particles thickness at the CL (W = 3.35 ± 0.28 nm, Fig. 5C) is significantly higher compared to the surroundings, both inside the corresponding osteon (W = 3.22 ± 0.22 nm, p < 0.01) as well as outside (W = 3.33 ± 0.18 nm, p = 0.048). Considering particle length (Fig. 5D), all three distributions are asymmetric with tails extending towards larger values. The CL distribution has two peaks and exhibits shorter crystals (L = 23.73 ± 8.14 nm) than the osteon (L = 25.88 ± 7.57 nm, p < 0.01) and the outer bone (L = 26.43 ± 8.22 nm, p < 0.01). Mineral particles at the CL seem to be slightly more oriented (ρ = 0.34 ± 0.1, Fig. 5E) than those inside the osteon (ρ = 0.32 ± 0.11, p < 0.01) and outside (ρ = 0.33 ± 0.1, p < 0.01). The lattice spacing (Fig. 5F) is lower at the CL (c-lattice = 6.884 ± 0.003 Å) compared to the other two regions (osteon: c-lattice = 6.884 ± 0.003 Å, p < 0.01; outer bone: c-lattice = 6.885 ± 0.003 Å, p < 0.01). Finally, zinc intensity is significantly higher in the CL than in the other two bone regions (Fig. 5A).

**Fig. 5:**
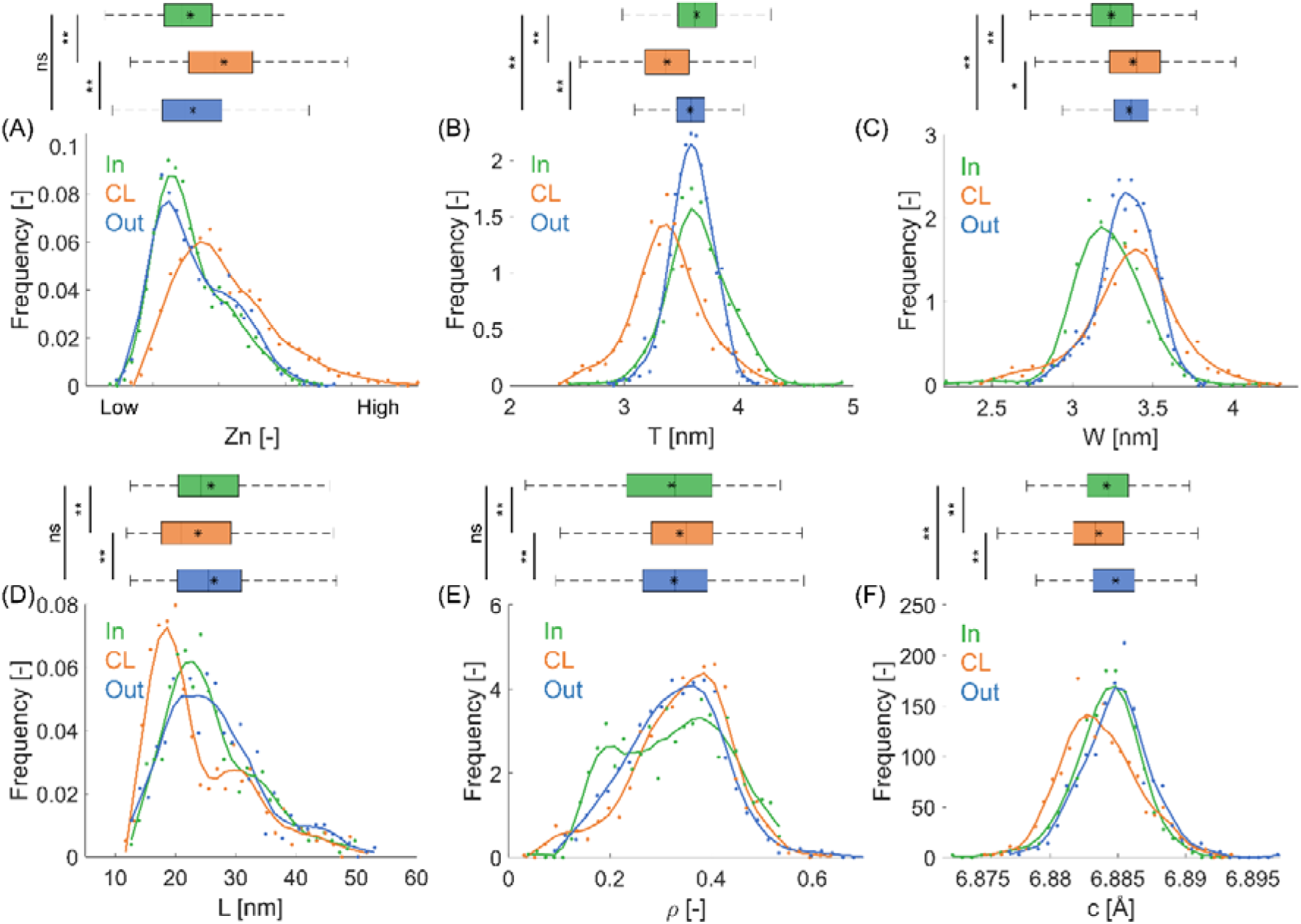
Frequency distributions and box plots of mineral particles properties: (A) zinc intensity Zn, (B) uncorrected particle thickness T, (C) mineral content corrected particle thickness W, (D) particle length L, (E) degree of alignment ρ, (F) lattice spacing c-lattice. All parameters are calculated in 3 rectangular regions (2 x 10 µm) including inner osteon (In), cement line (CL) and outer bone (Out). In and Out regions are 2 µm away from the CL. Original data represented as scattered points and smoothed data (with loess) as full lines. Boxes range from the 25^th^ to the 75^th^ percentiles, the median and mean values are indicated by a vertical line and an asterisk, respectively. Whiskers show the data range from the 5^th^ to the 95^th^ percentile. Statistical significance between the data sets is indicated as follows: *p < 0.05, **p < 0.01, and ns (not significant).

Despite the strong heterogeneity of the analyzed regions, which is in line with earlier investigations of osteonal bone^27, 62^, we could show that the CL features shorter but thicker mineral particles and has a slightly contracted c axis in comparison to the surrounding bone. Our finding on particle length is consistent with a previous work analyzing a portion of a single osteon with X-ray scattering tensor tomography.^27^ Considering the T parameter, similarly to Grünewald et al.,^27^ we found smaller values at the CL. However, when correcting the T parameter using qBEI data to account for differences in mineral volume fraction observed across the CL, we obtained the more reliable W parameter^63, 64^, indicating that the CL has thicker mineral particles than adjacent tissue. The geometrical dimensions of the mineral particles depend on the mineralization process and change during mineralization^64^. Rapidly deposited woven bone has thicker and shorter particles^65^, partly because in less ordered bone the particles should have more space to grow in thickness. Similar to woven bone, the CL is formed quicker and mineralizes faster than lamellar bone.^26^ In addition to the arrangement of collagen fibrils, other factors that can influence particle dimensions are matrix composition and water content.^66^ The CL is rich in non-collagenous proteins such as osteocalcin and osteopontin,^21, 23, 25, 67^ which have the ability to regulate crystal growth, size and shape.^68^ The CL also contains lower amount of collagen compared to the surrounding bone matrix,^24^ which may influence mineral-collagen packing. One additional specificity of the CL is the scarcity of nanochannels, which are, on the other hand, present in lamellar bone.^69^ The accumulation of thicker mineral particles in the CL may lead to the closure of the nanochannels.

A slightly shorter c axis of mineral crystal in the CL agrees with previous work ^27^: it may be due to the presence of trace elements like zinc^57^ or it may indicate a lower degree of carbonate substitution, typically occurring during bone maturation.^70^ Indeed, decreasing c-lattice with increasing zinc content has been reported during formation of hydroxyapatite in vitro.^71^ Zinc may be incorporated in the course of CL mineralization, which happens in proximity with blood vessels. Although the CL has a different mineralization process than lamellar bone, with slower secondary mineralization^26^, additional analysis for example based on Fourier Transform Infrared Spectroscopy (FTIR)^72^ would be required to characterize the amount of carbonate substitution in the CL. One additional feature responsible the contraction of the c axis could be compressive strains in the mineral induced by the mineralization process^73, 74^: the higher mineral content of the CL may impose larger stresses on the apatite crystals.

The organization of mineralized collagen fibrils in the CL is still debated. One study based on FIB-SEM^55^ indicates that CL fibrils are oriented in three or more predominant directions and have higher directional dispersion than in lamellar bone. The degree of alignment quantified here with the ρ parameter shows great spatial variability across the different regions (Fig. 4F and Supplementary Information, Fig. S11). This is also reflected in the frequency distributions of ρ: the CL tends to have a marginally higher degree of alignment than immediately adjacent bone (Fig. 5E), but no differences are detected when analyzing a larger region (Supplementary Information, Fig. S11).

By correlating mineral content and mechanical properties, we found that the CL requires a higher mineral content to achieve the same stiffness and hardness as the surrounding bone (Fig. 3B). We searched for a possible explanation by focusing on the mineral characteristics, and we observed that mineral particles at the CL appear thicker and shorter compared to nearby tissue. These findings can be interpreted with a mechanical shear-lag model (Supplementary Note 1).^11^ Assuming a staggered arrangement of the mineral particles into the organic matrix (Fig. 6A), the model allows to compute the elastic modulus and the tensile strength of the resulting composite as a function of mineral volume fraction and mineral particle dimensions.^3, 42^ Introducing the aspect ratio of the mineral particles as length over thickness, we obtain an average value of about 8 for the lamellar bone and 7 for the CL. Mechanically, a low aspect ratio is clearly reflected in lower stiffness and tensile strength for the same mineral volume fraction (Figs. 6B and C). This suggests that mineral particles in the CL are less effective at reinforcing the organic matrix. Previous works have highlighted that a higher aspect ratio is beneficial for nanoscale stiffness thanks to an increased stress transfer along the crystal length.^75, 76^ Longer mineral particles should also provide a higher surface area for mineral-collagen interactions^77^ and restrict collagen mobility.^78^ In contrast, shorter crystals may leave a greater number of unreinforced regions in the collagen network, leading to higher post-yield deformability.^79^

**Figure 6:**
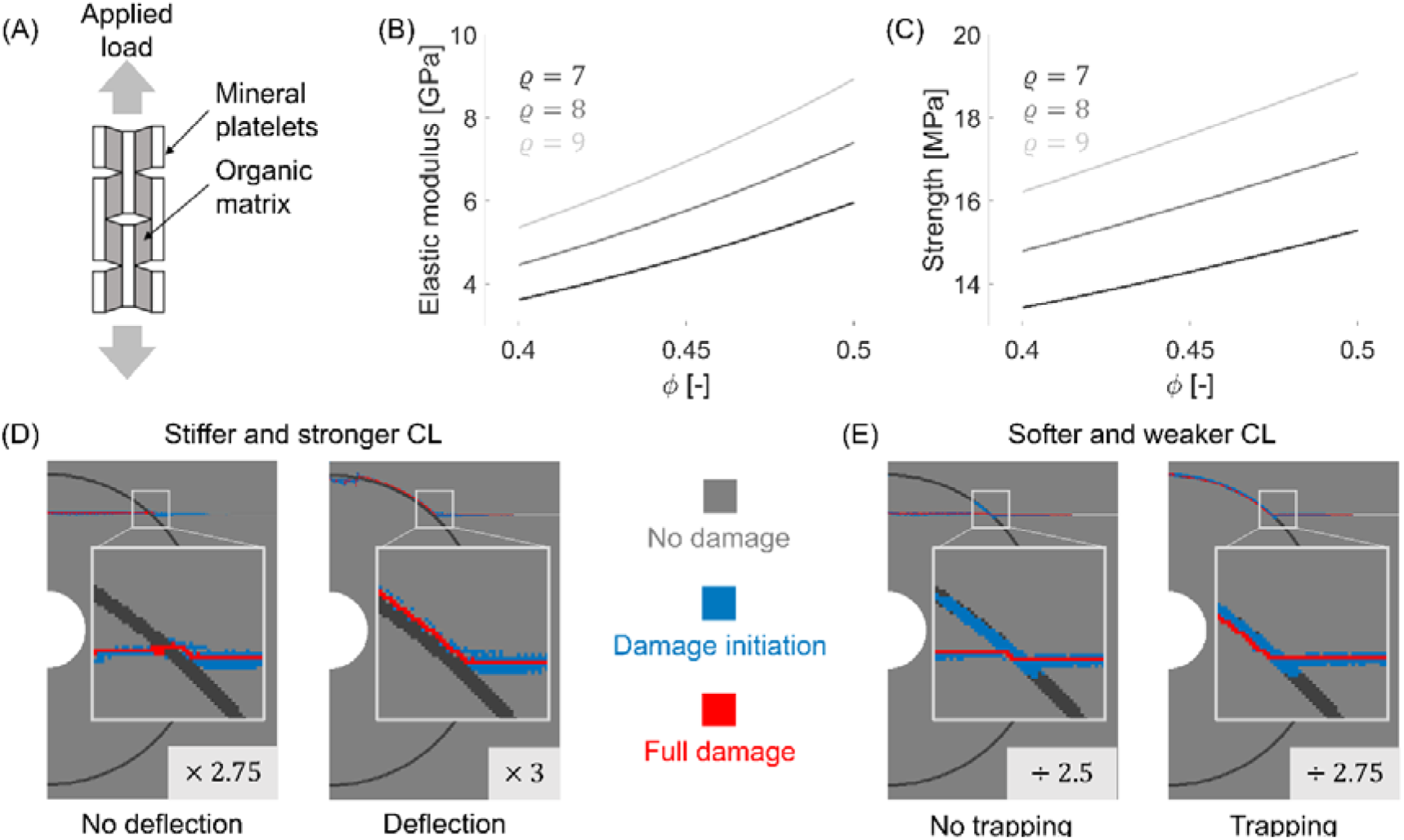
(A) Schematic representation of the staggered composite model which assumes mineral particles having plate-like shape embedded into a ductile organic matrix under tensile loading. Theoretical predictions of (B) the composite elastic modulus and (C) tensile strength as a function of mineral volume fraction, for different aspect ratios of the mineral particles. Simulations of damage propagation considering a half-osteon bordered by a CL and embedded into interstitial bone, considering (D) a stiffer/stronger but more brittle CL and (E) a softer/weaker but tougher CL compared to adjacent bone. Damage patterns are color-coded: blue indicates damage initiation and red indicates full damage. The numbers at the bottom of the images indicated the corresponding elastic contrast (i.e. ratio between properties of CL and bone).

### On the protective role of the cement line

Finally, we use a proof-of-concept damage-based computational model^80^ to challenge the assumed ability of the CL to protect the osteon (Supplementary Note 2). We choose a simplified scenario featuring half an osteon, with a Haversian canal, bordered by a CL and embedded into interstitial bone (Fig. 6D). We define the mechanical contrast as the ratio between the stiffness (and the strength) of the CL compared to those of the surrounding bone. We assumed that a stiffer (stronger) CL is also more brittle (lower fracture strain); likewise, a softer (weaker) CL is also tougher (higher fracture strain) than the osteon (Fig. S12). Our simulations indicate that a stiff, strong and brittle CL can deflect damage only if the differences in mechanical properties with respect to the osteon are large: a mechanical contrast of more than 300% is required (Fig. 6D), which is significantly greater than the 20% difference measured experimentally in this work. A soft, weak and tough CL may also protect the osteon by trapping damage, but again the required decrease in properties for full trapping is substantial, i.e. a factor of 275% (Fig. 6E). Several previous computational works have explored the role of the CL in the damaging behavior of osteonal bone. Considering the uncertainties on CL properties, both weaker and stiffer CL have been assumed.^35–38, 81–83^ Our results support the view of a stiffer, stronger and most likely more brittle CL but also highlight that the required differences in mechanical properties between CL and surrounding bone for effective damage protection may be substantial. Although the mechanical contrast is expected to change when considering different notch lengths and positions, we still suspect that the CL alone may be of limited help for the mechanical protection of the osteon, perhaps hampering only the propagation of short cracks meeting the CL with a favorable trajectory (i.e. at small angles). Yet, experimental observations often reports cracks to propagate along CL.^30, 37, 84^ Clearly, protecting the osteon and the blood vessels hosted in the Haversian canal is a critical task that cannot be solved by the CL alone. Other features should play an important role, such as pre-stresses^73, 74^ and lamellar arrangements. Indeed, the rotated plywood lamellar architecture around the osteon has been demonstrated to be a very effective way to improve strength, toughness and damage tolerance.^14, 80^ Lamellar bone also occupies a considerable portion of the osteon, much greater than the space reserved to the thin CL. Indeed, lamellar architectures are not limited to bone but this is a universal strategy of nature to build damage tolerant mineral-reinforced materials.^3, 85–87^

### Limitations

Some limitations of this study should be acknowledged. First, the number of samples and osteons had to be restricted due to technical and time constraints of our multimodal analysis (e.g., limited availability of synchrotron beamtime). Like many studies focusing on micro- and nanoscopic material properties,^41, 48, 52, 62, 69, 88^ we prioritized the detailed characterization of a limited number of samples rather than broader, less in-depth testing of many samples. This strategy is supported by the strong spatial heterogeneity of the micro- and nanostructure of bone, which requires a mapping of large regions to obtain robust results. Despite this constraint, all measured compositional, structural, and mechanical properties were consistent across all analyzed regions. To further increase the robustness of our findings, future research should include larger and more diverse sample groups, incorporating variations in sex, age, and anatomical location. Second, bone samples were tested in a dehydrated state to enable measurements of the mineral content and a correlative analysis with the mechanical properties. It is known that ethanol removes water while maintaining hydrogen bonding with collagen.^89^ As a consequence, indentation modulus and hardness in dehydrated samples tend to be higher compared to hydrated conditions.^50, 90^ However, our interest in only relative differences between CL and neighboring bone rather than in absolute values mitigates the impact of this limitation. Third, a pixel-by-pixel superimposition of measurements of mechanical (nIND) and mineral (SAXS/WAXS) properties was not possible due sample preparation (see Methods) and corresponding slightly different analyzed sections (about 15 µm apart). As a result, only qualitative correlations could be established. However, the analyzed regions still display comparable structural features, such as Haversian canals and CL. These features were identifiable in both fluorescence (zinc) and qBEI maps, allowing for manual image registration and alignment. qBEI was performed at a relatively high resolution to have enough pixels inside the CL while minimizing partial volume effects.^91^ Although at such resolution the qBEI signal should be only marginally influenced by additional contrast coming from the lamellar arrangement, we did see small mineral differences at the level of bone lamellae having different orientations but we cannot say how much of this variation in mineral content is due to orientation effects. This is much less problematic at the CL, considering the higher values of mineral content measured there. Orientation effects are also present in SAXS/WAXS measurements with L and ρ being the most affected parameters. An accurate quantification of these effects would require systematic sample rotation^14^, which was not feasible within the constraints of the available synchrotron beamtime and is beyond the scope of the current study, warranting further investigation in future works.

## Conclusion

We demonstrated that the CL is stiffer and harder than the adjacent osteonal tissue challenging the widespread assumption that it acts as a soft, compliant interface. The central finding of our investigation is that the CL requires more mineral to attain the same stiffness and hardness as surrounding bone. We look for a possible explanation by investigating the nanoscale structure of the mineral particles, which are thicker but shorter in the CL. With the help of a micromechanical model we showed that mineral particles with a lower aspect ratio are less effective at reinforcing the collagen matrix. Using damage-based computer simulations, we questioned the assumed role of the CL to deviate cracks as the mechanical properties required for effectively performing such a task would be much different than those measured in our study.

## Methods

### Sample preparation

Femurs from two male donors, aged 40 and 81, with no known bone conditions, were acquired from the Department of Forensic Medicine of the Medical University of Vienna. This study complied with institutional ethical guidelines (EK no. 1757/2013). Immediately after extraction, the femurs were frozen at -20 °C for preservation. Femoral shafts far from the metaphysis were cut transversally and sectioned into 6 cubic samples (3 per femur), each approximately 1.5 cm in size, using a water-cooled wire diamond saw (WireTec, Germany). Following this, the samples underwent dehydration and embedding in poly-methyl methacrylate (PMMA).^69, 92^ The upper transverse surfaces were then ground with sandpapers of progressively finer grit sizes (P1200, P2400) and polished using a diamond suspension down to 1 µm particle size on a silk cloth under glycol irrigation (Logitech PM5, UK), to minimize microcrack formation during preparation.^93^ Samples were then characterized with quantitative backscattered electron imaging (qBEI), nanoindentation (nIND), scanning electron microscopy (SEM) and small- and wide-angle X-ray scattering. An overview table summarizing which measurements were made at which samples and at which osteons is given in the Supplementary Information, Table S1.

### Quantitative backscattered electron imaging

We used quantitative backscattered electron imaging (qBEI) to measure the local mineral content of the CL and surrounding bone. This technique enables a quantitative two-dimensional assessment of the calcium content with a spatial resolution of less than 1 µm, as required by the tiny dimensions of the CL. Briefly, qBEI counts the electrons which are backscattered from the sample surface during scanning electron microscopy (SEM). The backscattered-electron signal is calibrated with carbon and aluminum standards to convert grey level images into spatial maps of the calcium content (in Ca wt %), according to a well-established procedure.^91, 92, 94, 95^ Prior to qBEI experiments, the samples were coated with a thin carbon layer using vacuum evaporation (Agar SEM carbon coater, Agar, Stansted, UK) to achieve a conductive surface. qBEI measurements were performed with a SEM equipped with a zirconium-coated field emission cathode (Zeiss SEM SUPRA 40, Germany). The SEM operated at 20 kV, with a working distance of 10 mm and a probe current ranging between 280 and 320 pA. Initial overview scans were done at a lower resolution (1.76 µm/pixel) with a wider field of view to identify osteons with a visible CL in different locations. A total of 12 osteons of various dimensions and mineral content were preselected and analyzed at higher magnification (isotropic pixel size of 0.57 μm and field of view of 1024 × 768 pixels).

### Second harmonic generation imaging

To qualitatively assess the lamellar organization of the selected osteons and surrounding bone, second harmonic generation (SHG) imaging was conducted on the same regions previously measured by qBEI. Imaging was performed using a confocal laser scanning microscope (Leica TCS SP8 DLS, Germany) equipped with a 40× oil immersion objective (HC PL APO 40×/1.30 OIL). A pulsed infrared laser (Spectra-Physics, Milpitas, US) operating at 910 nm was utilized, with backward signal collected at wavelengths between 450 and 460 nm. Images were captured with a nominal pixel size of 379 nm across a 1024 × 1024 pixels field of view, scanned at a rate of 400 Hz. Multiple regions were imaged and subsequently stitched together with an overlapping of 10% using the Leica Application Suite X.

### Nanoindentation

To measure the local mechanical properties, we used nanoindentation (nIND) on regions identified and characterized with qBEI (Fig. 1B). To map the mechanical properties with a similar resolution as used for measuring the mineral content, we imposed a maximum load of 500 µN, corresponding to a penetration depth of about 150 nm. This depth allows a spacing between adjacent indents of 1 µm, which should be roughly 7 times the contact depth to avoid overlapping of the inelastic deformation fields.^96^ Nanoindentation was performed with a Triboindenter TI 950 (Bruker, US) equipped with a Berkovich diamond tip (100 nm tip radius). A force-controlled trapezoidal load function was employed, involving loading for 8 seconds, holding for 20 seconds and unloading for 8 seconds. Prior to indentation, sample surfaces were scanned with the tip at 2 µN contact force over regions of 50 × 50 µm to measure the average surface roughness, which was 7.97 ± 10.47 nm. As the indentation depth was about ten times the surface roughness, the impact of roughness can be neglected.^97, 98^ Several indentation grids with a constant spacing of 1 µm were performed on a subset of the osteons previously analyzed with qBEI. We selected 9 regions belonging to 8 different osteons, where the CLs were most visible and had the largest width, resulting in 4966 indents. Before indentation, the probe contact area was calibrated on fused quartz. Force-depth curves were analyzed with the well-established Oliver-Pharr method,^99^ to obtain indentation hardness (H) and reduced modulus (E_r_).

### High-resolution backscattered and secondary electron imaging

To locate the position of the indents with respect to the CLs, the indented regions were rescanned using the same SEM as used for qBEI (Zeiss SEM Supra 40) at higher magnification, yielding an isotropic pixel size of approximately 76 nm. The high-resolution images were captured in two modalities simultaneously: secondary electron imaging (SEI) was used to map the surface topography allowing to detect the tiny traces of the indents caused by permanent deformation (Fig. 1C); backscattered electron imaging (BEI) facilitated the distinction between CLs (appearing brighter) and the surrounding bone based on different grey values (atomic number contrast), and allowed to distinguish layers with different textures, reflecting the lamellar structure of osteonal bone (Fig. 1D). Here, we used the terminology of bright and dark lamellae referring to the bright and dark layers visible in polarized light^100^ or SHG (Supplementary Information, Fig S1). By overlaying the indent traces (visible in SEI) on the high resolution BE images of the bone surface, we could classify the indents as belonging to the CLs, the dark or bright lamellae (of the same osteon of the CL) or the surrounding older bone outside the osteon (Fig. 1E). Indents located within cracks or pores (e.g. osteocyte lacunae) were excluded from the analysis. We also neglected indents only partially falling into the CLs (red symbols in Fig. 1E).

### Correlation between mineral content and mechanical properties

To extract the mineral content at the very same positions probed with nanoindentation, quantitative maps of the mineral content obtained with qBEI were superimposed with nanoindentation grids, exploiting high resolution qualitative BEI/SEI images showing the CL and the indent traces, respectively (Fig 1E). Using a custom image registration technique implemented in MATLAB, five to six landmarks were manually selected in both qBEI and BEI/SEI images. These landmarks included for example the porosities, and were used to compute a similarity transformation, which preserves angles and aspect ratios while allowing for translation, rotation, and uniform scaling, using the fitgeotrans function from MATLAB Image Processing Toolbox (R2022a, The MathWorks, Natick, MA, USA). Indent coordinates could be mapped to the qBEI reference frame using the obtained transformation matrix. For the correlation with the mechanical properties, the mineral content was averaged over a 3 × 3 pixels region centered around each indent.

### Small- and wide-angle X-ray scattering

After assessing the mineral content (qBEI), the fiber organization (SHG) and the local mechanical properties (nIND), the same samples were examined with scanning small- and wide-angle X-ray scattering (SAXS and WAXS) to measure mineral particle properties, with sub-micrometer resolution. The samples were sectioned into very thin slices (Fig. 1F), approximately 2-3 µm thick, using a microtome (Leica SM2500, Leica, Nussloch, Germany). The thin sections were mounted on a silicon nitride X-ray transparent window (Norcada Inc., Canada) and fixed to custom made aluminum frames for SAXS/WAXS experiments (Supplementary Information, Fig. S2). All regions to be analyzed were preselected with qBEI and, prior to X-ray scattering experiments, the slices were imaged with an optical microscope (Keyence VHX Microscope, Belgium) to locate the positions of the pre-selected regions. Scanning X-ray scattering was performed at the microfocus beamline (ID13) of the European Synchrotron Radiation Facility (ESRF, Grenoble, France). A monochromatic beam with an energy of 15.2 keV and focal spot size of 0.5 µm was used. SAXS and WAXS patterns were recorded simultaneously using a Eiger 4M detector (DECTRIS, Baden-Daettwill, Switzerland) with a pixel size of 75 × 75 µm. The sample-to-detector distance was 237.25 mm, calibrated with an NIST aluminum oxide standard. Each measurement point was illuminated for 20 ms. Parallel to the scattering measurements, quantitative X-ray fluorescence (XRF) maps were recorded using an XRF detector placed approximately 20 mm away from the sample. A total of 22 regions having typical dimensions of 200 x 200 µm^2^, belonging to 20 different osteons were mapped with an isotropic pixel size of 0.5 µm. The SAXS and WAXS data were processed with the software named DPDAK (Directly Programmable Data Analysis Kit).101 After background correction, the SAXS and WAXS profiles have been integrated over the azimuthal angle to obtain radial intensity profiles I(q), with q being the length of the scattering vector **q**.^63,102^ The radial intensity profile has been used to obtain information on mineral.^46^ Assuming that mineral crystals exhibit a thin platelet morphology, the so-called T parameter (Figs. 1G-H) is defined as volume-to-surface ratio of the particles 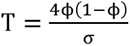, with ϕ representing the mineral volume fraction and σ denoting the platelet surface per unit volume.^103^ The T parameters is a good approximation for the average particle thickness if the mineral volume fraction in the interaction volume is close to 0.5 and assuming the thickness as smallest dimension of the particles a « b ≤ c.^58^ As in our study we compared regions with large differences in mineral content, including lowly mineralized osteons and highly mineralized CLs^26^, we also calculated the W parameter, which is a more precise description of the T parameter obtained by correction for variation in □ based on local measures of the mineral content.^58^ The W parameter is defined as 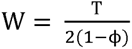, with ϕ being now the spatially varying volume fraction, which is measured with qBEI on approximately the same locations characterized by X-ray scattering. To assess the platelet length L along the c axis (Figs. 1G-H), the full width at half maximum (FWHM) of the (002) peak intensity in the WAXS signal is used. Specifically, we applied Scherrer’s equation: 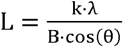, where λ is the wavelength, θ is the Bragg angle, k is the Scherrer constant (related to crystallite shape), and B is the FWHM of the peak.^104^ The position of the (002) peak indicates the lattice spacing (i.e. the space between atoms) in the crystallographic (002) direction. This is used to compute the c-lattice constant (Fig. 1G-H), which informs on possible distortions of the HAP crystal, for example due to trace elements.^105^ Finally, we derived the degree of orientation, denoted as ρ parameter, by integrating the azimuthal SAXS intensity profile.^63^ The degree of orientation describes how well the mineral platelets are aligned along a common direction within the plane perpendicular to the incident X-ray beam: ρ value of 1 indicates a uniform orientation, while ρ = 0 represents a completely random orientation (Fig. 1G-H).^106^

Pixels with insufficient scattering signal to derive mineral crystal parameters were excluded from the analysis by setting thresholds for the total scatter intensity.^64^ To locate the CL in the position resolved maps of the crystal parameters, we used XRF images acquired alongside the X-ray scattering. As calcium maps were heavily influenced by surface waviness, we relied on zinc maps, which provided a good contrast between the CL and the surrounding bone. Indeed, it is known that zinc and other heavy metals accumulate in the CL.^57^ The identification of the CL was further verified by overlaying zinc and qBEI maps, in the latter the CL appears brighter^26^ (Supplementary Information, Fig. S3).

Mineral parameters were analyzed in the proximity of the CL and plotted as a function of the distance to the CL (Fig. 4). In the distance plots, three regions were identified: the CL, the corresponding inner osteon (In), and the surrounding outer bone (Out). The CL was defined as a 2-µm-thick layer centered around the peak in the Zn signal. Due to the heterogeneity of mineral properties, we analyzed only zones in the vicinity of the CL: inner and outer regions extended up to 8 µm away from the CL. To compare mineral properties, we calculated frequency distributions of mineral parameters in the CL and in two additional regions having the same width of the CL and located ±2 µm from the CL, thus falling inside and outside the osteon, respectively (Fig. 5). To enhance readability, all frequency distributions were smoothed using a local regression with weighted linear least squares and a first-degree polynomial model (loess fit). Overall, we measured 22 locations belonging to 20 different osteons. Six locations were discarded due to technical issues and ten locations due to sample artifacts (waviness and cracks), which cannot be avoided when preparing large and thin sections as required for synchrotron experiments at sub-micrometer resolution. In the remaining dataset, we analyzed a total of 6 osteons (Supplementary Information, Fig. S3) with 18 distance plots (3 for each osteon), corresponding to a bone surface of 3600 µm^2^.

### Statistical analysis

We compared differences in mechanical properties and mineral properties among the three regions (CL, In, and Out) by performing a Kruskal-Wallis test, chosen due to the non-normal distribution of the unpaired data. When significant differences were detected, pairwise comparisons between locations were conducted using post-hoc tests with Bonferroni correction for multiple comparisons minimizing false positive results. We examined the strength of the relationships between the mean calcium content of osteon, CL and different regions using Spearman’s rank correlation, selected for its suitability with non-normal distribution of the data. Significance levels were set at p < 0.05, and all p-values were two-sided. All statistical analyses were performed in MATLAB 2022a.

## Acknowledgments

The authors thank Alexandra Tits and Sarvesh Bhogaokar for their help during the beamtime and acknowledge the European Synchrotron Radiation Facility (ESRF) in Grenoble, France for granting beamtime under the proposal SC-5337. They also thank Phaedra Messmer, Petra Keplinger, and Sonja Lueger from LBIO for the excellent sample preparation. AC is supported by the French Community of Belgium as part of a FRIA (Fund for Research Training in Industry and Agriculture) grant (n°1E03621F) and by the FWO and F.R.S.-FNRS under the Excellence of Science (EOS) program (EOS No. 40007553). MH and SB gratefully acknowledge financial support from the Austrian Social Health Insurance Fund (OEGK) and the Austrian Workers’ Compensation Board (AUVA). MR and RW thank the Max Planck Queensland Centre for the Materials Science of Extracellular Matrices for support. Additionally, this work was supported by the German Research Foundation grant SFB1444-P7 to VS.

## References

(1) Currey, J. D. Bones: Structure and Mechanics; Princeton University Press, 2002.

(2) Bergmann, G.; Deuretzbacher, G.; Heller, M.; Graichen, F.; Rohlmann, A.; Strauss, J.; Duda, G. N. Hip Contact Forces and Gait Patterns from Routine Activities. J. Biomech. 2001, 34 (7), 859–871. 10.1016/S0021-9290(01)00040-9.

(3) Fratzl, P.; Weinkamer, R. Nature’s Hierarchical Materials. Prog. Mater. Sci. 2007, 52 (8), 1263–1334. 10.1016/j.pmatsci.2007.06.001.

(4) Studart, A. R. Biological and Bioinspired Composites with Spatially Tunable Heterogeneous Architectures. Adv. Funct. Mater. 2013, 23 (36), 4423–4436. 10.1002/adfm.201300340.

(5) Wagermaier, W.; Klaushofer, K.; Fratzl, P. Fragility of Bone Material Controlled by Internal Interfaces. Calcif. Tissue Int. 2015, 97 (3), 201–212. 10.1007/s00223-015-9978-4.

(6) Zimmermann, E. A.; Ritchie, R. O. Bone as a Structural Material. Adv. Healthc. Mater. 2015, 4 (9), 1287–1304. 10.1002/adhm.201500070.

(7) Weinkamer, R.; Fratzl, P. Solving Conflicting Functional Requirements by Hierarchical Structuring—Examples from Biological Materials. MRS Bull. 2016, 41 (09), 667–671. 10.1557/mrs.2016.168.

(8) Dunlop, J. W. C.; Weinkamer, R.; Fratzl, P. Artful Interfaces within Biological Materials. Mater. Today 2011, 14 (3), 70–78. 10.1016/S1369-7021(11)70056-6.

(9) Beniash, E. Biominerals—Hierarchical Nanocomposites: The Example of Bone. WIREs Nanomedicine Nanobiotechnology 2011, 3 (1), 47–69. 10.1002/wnan.105.

(10) Gupta, H. S.; Seto, J.; Wagermaier, W.; Zaslansky, P.; Boesecke, P.; Fratzl, P. Cooperative Deformation of Mineral and Collagen in Bone at the Nanoscale. Proc. Natl. Acad. Sci. 2006, 103 (47), 17741–17746. 10.1073/pnas.0604237103.

(11) Jäger, I.; Fratzl, P. Mineralized Collagen Fibrils: A Mechanical Model with a Staggered Arrangement of Mineral Particles. Biophys. J. 2000, 79 (4), 1737–1746. 10.1016/S0006-3495(00)76426-5.

(12) Fantner, G. E.; Hassenkam, T.; Kindt, J. H.; Weaver, J. C.; Birkedal, H.; Pechenik, L.; Cutroni, J. A.; Cidade, G. A. G.; Stucky, G. D.; Morse, D. E.; Hansma, P. K. Sacrificial Bonds and Hidden Length Dissipate Energy as Mineralized Fibrils Separate during Bone Fracture. Nat. Mater. 2005, 4 (8), 612–616. 10.1038/nmat1428.

(13) Weiner, S.; Wagner, H. D. THE MATERIAL BONE: Structure-Mechanical Function Relations. Annu. Rev. Mater. Sci. 1998, 28 (1), 271–298. 10.1146/annurev.matsci.28.1.271.

(14) Wagermaier, W.; S. Gupta, H.; Gourrier, A.; Burghammer, M.; Roschger, P.; Fratzl, P. Spiral Twisting of Fiber Orientation inside Bone Lamellae. Biointerphases 2006, 1 (1), 1–5. 10.1116/1.2178386.

(15) Weiner, S.; Arad, T.; Sabanay, I.; Traub, W. Rotated Plywood Structure of Primary Lamellar Bone in the Rat: Orientations of the Collagen Fibril Arrays. Bone 1997, 20 (6), 509–514. 10.1016/S8756-3282(97)00053-7.

(16) Fratzl, P.; Gupta, H. S.; Fischer, F. D.; Kolednik, O. Hindered Crack Propagation in Materials with Periodically Varying Young’s Modulus—Lessons from Biological Materials. Adv. Mater. 2007, 19 (18), 2657–2661. 10.1002/adma.200602394.

(17) Peterlik, H.; Roschger, P.; Klaushofer, K.; Fratzl, P. From Brittle to Ductile Fracture of Bone. Nat. Mater. 2006, 5 (1), 52–55. 10.1038/nmat1545.

(18) Weiner, S.; Traub, W.; Wagner, H. D. Lamellar Bone: Structure–Function Relations. J. Struct. Biol. 1999, 126 (3), 241–255. 10.1006/jsbi.1999.4107.

(19) Shahar, R.; Weiner, S. Open Questions on the 3D Structures of Collagen Containing Vertebrate Mineralized Tissues: A Perspective. J. Struct. Biol. 2018, 201 (3), 187–198. 10.1016/j.jsb.2017.11.008.

(20) McKEE, M. D.; Nanci, A. Osteopontin and the Bone Remodeling Sequence: Colloidal-Gold Immunocytochemistry of an Interfacial Extracellular Matrix Protein ^a^. Ann. N. Y. Acad. Sci. 1995, 760 (1), 177–189. 10.1111/j.1749-6632.1995.tb44629.x.

(21) Lassen, N. E.; Andersen, T. L.; Pløen, G. G.; Søe, K.; Hauge, E. M.; Harving, S.; Eschen, G. E. T.; Delaisse, J. Coupling of Bone Resorption and Formation in Real Time: New Knowledge Gained From Human Haversian BMUs. J. Bone Miner. Res. 2017, 32 (7), 1395–1405. 10.1002/jbmr.3091.

(22) Sims, N. A. Osteoclast-Derived Coupling Factors: Origins and State-of-Play Louis V Avioli Lecture, ASBMR 2023. J. Bone Miner. Res. 2024, 39 (10), 1377–1385. 10.1093/jbmr/zjae110.

(23) McKee, M. D.; Nanci, A. Osteopontin at Mineralized Tissue Interfaces in Bone, Teeth, and Osseointegrated Implants: Ultrastructural Distribution and Implications for Mineralized Tissue Formation, Turnover, and Repair. Microsc. Res. Tech. 1996, 33 (2), 141–164. 10.1002/(SICI)1097-0029(19960201)33:2<141::AID-JEMT5>3.0.CO;2-W.

(24) Milovanovic, P.; vom Scheidt, A.; Mletzko, K.; Sarau, G.; Püschel, K.; Djuric, M.; Amling, M.; Christiansen, S.; Busse, B. Bone Tissue Aging Affects Mineralization of Cement Lines. Bone 2018, 110, 187–193. 10.1016/j.bone.2018.02.004.

(25) Skedros, J. G.; Holmes, J. L.; Vajda, E. G.; Bloebaum, R. D. Cement Lines of Secondary Osteons in Human Bone Are Not Mineral-Deficient: New Data in a Historical Perspective. Anat. Rec. A. Discov. Mol. Cell. Evol. Biol. 2005, 286A (1), 781–803. 10.1002/ar.a.20214.

(26) Cantamessa, A.; Blouin, S.; Rummler, M.; Berzlanovich, A.; Weinkamer, R.; Hartmann, M. A.; Ruffoni, D. The Mineralization of Osteonal Cement Line Depends on Where the Osteon Is Formed. JBMR Plus 2025, ziaf114. 10.1093/jbmrpl/ziaf114.

(27) Grünewald, T. A.; Johannes, A.; Wittig, N. K.; Palle, J.; Rack, A.; Burghammer, M.; Birkedal, H. Bone Mineral Properties and 3D Orientation of Human Lamellar Bone around Cement Lines and the Haversian System. IUCrJ 2023, 10 (2), 189–198. 10.1107/S2052252523000866.

(28) Kerschnitzki, M.; Wagermaier, W.; Roschger, P.; Seto, J.; Shahar, R.; Duda, G. N.; Mundlos, S.; Fratzl, P. The Organization of the Osteocyte Network Mirrors the Extracellular Matrix Orientation in Bone. J. Struct. Biol. 2011, 173 (2), 303–311. 10.1016/j.jsb.2010.11.014.

(29) Nalla, R. K.; Kruzic, J. J.; Kinney, J. H.; Ritchie, R. O. Mechanistic Aspects of Fracture and R-Curve Behavior in Human Cortical Bone. Biomaterials 2005, 26 (2), 217–231. 10.1016/j.biomaterials.2004.02.017.

(30) Mohsin, S.; O’Brien, F. J.; Lee, T. C. Osteonal Crack Barriers in Ovine Compact Bone. J. Anat. 2006, 208 (1), 81–89. 10.1111/j.1469-7580.2006.00509.x.

(31) Koester, K. J.; Ager, J. W.; Ritchie, R. O. The True Toughness of Human Cortical Bone Measured with Realistically Short Cracks. Nat. Mater. 2008, 7 (8), 672–677. 10.1038/nmat2221.

(32) Qiu, S.; Sudhaker Rao, D.; Fyhrie, D. P.; Palnitkar, S.; Parfitt, A. M. The Morphological Association between Microcracks and Osteocyte Lacunae in Human Cortical Bone. Bone 2005, 37 (1), 10–15. 10.1016/j.bone.2005.01.023.

(33) Gauthier, R.; Follet, H.; Olivier, C.; Mitton, D.; Peyrin, F. 3D Analysis of the Osteonal and Interstitial Tissue in Human Radii Cortical Bone. Bone 2019, 127, 526–536. 10.1016/j.bone.2019.07.028.

(34) Hull, D.; Clyne, T. W. An Introduction to Composite Materials, 2nd ed.; Cambridge Solid State Science Series; Cambridge University Press: Cambridge, 1996. 10.1017/CBO9781139170130.

(35) Vergani, L.; Colombo, C.; Libonati, F. Crack Propagation in Cortical Bone: A Numerical Study. Procedia Mater. Sci. 2014, 3, 1524–1529. 10.1016/j.mspro.2014.06.246.

(36) Gustafsson, A.; Wallin, M.; Khayyeri, H.; Isaksson, H. Crack Propagation in Cortical Bone Is Affected by the Characteristics of the Cement Line: A Parameter Study Using an XFEM Interface Damage Model. Biomech. Model. Mechanobiol. 2019, 18 (4), 1247–1261. 10.1007/s10237-019-01142-4.

(37) Giner, E.; Belda, R.; Arango, C.; Vercher-Martínez, A.; Tarancón, J. E.; Fuenmayor, F. J. Calculation of the Critical Energy Release Rate G c of the Cement Line in Cortical Bone Combining Experimental Tests and Finite Element Models. Eng. Fract. Mech. 2017, 184, 168–182. 10.1016/j.engfracmech.2017.08.026.

(38) Nobakhti, S.; Limbert, G.; Thurner, P. J. Cement Lines and Interlamellar Areas in Compact Bone as Strain Amplifiers – Contributors to Elasticity, Fracture Toughness and Mechanotransduction. J. Mech. Behav. Biomed. Mater. 2014, 29, 235–251. 10.1016/j.jmbbm.2013.09.011.

(39) Montalbano, T.; Feng, G. Nanoindentation Characterization of the Cement Lines in Ovine and Bovine Femurs. J. Mater. Res. 2011, 26 (8), 1036–1041. 10.1557/jmr.2011.46.

(40) Zhou, Y.; Kastner, M. J.; Tighe, T. B.; Du, J. Elastic Modulus Mapping for Bovine Cortical Bone from Submillimeter-to Submicron-Scales Using PeakForce Tapping Atomic Force Microscopy. Extreme Mech. Lett. 2020, 41, 101031. 10.1016/j.eml.2020.101031.

(41) Gupta, H. S.; Schratter, S.; Tesch, W.; Roschger, P.; Berzlanovich, A.; Schoeberl, T.; Klaushofer, K.; Fratzl, P. Two Different Correlations between Nanoindentation Modulus and Mineral Content in the Bone–Cartilage Interface. J. Struct. Biol. 2005, 149 (2), 138–148. 10.1016/j.jsb.2004.10.010.

(42) Tits, A.; Blouin, S.; Rummler, M.; Kaux, J.-F.; Drion, P.; van Lenthe, G. H.; Weinkamer, R.; Hartmann, M. A.; Ruffoni, D. Structural and Functional Heterogeneity of Mineralized Fibrocartilage at the Achilles Tendon-Bone Insertion. Acta Biomater. 2023, 166, 409–418. 10.1016/j.actbio.2023.04.018.

(43) Reznikov, N.; Shahar, R.; Weiner, S. Three-Dimensional Structure of Human Lamellar Bone: The Presence of Two Different Materials and New Insights into the Hierarchical Organization. Bone 2014, 59, 93–104. 10.1016/j.bone.2013.10.023.

(44) Reznikov, N.; Shahar, R.; Weiner, S. Bone Hierarchical Structure in Three Dimensions. Biomineralization 2014, 10 (9), 3815–3826. 10.1016/j.actbio.2014.05.024.

(45) Hoo, R. P.; Fratzl, P.; Daniels, J. E.; Dunlop, J. W. C.; Honkimäki, V.; Hoffman, M. Cooperation of Length Scales and Orientations in the Deformation of Bovine Bone. Acta Biomater. 2011, 7 (7), 2943–2951. 10.1016/j.actbio.2011.02.017.

(46) Fratzl, P.; Gupta, H. S.; Paschalis, E. P.; Roschger, P. Structure and Mechanical Quality of the Collagen–Mineral Nano-Composite in Bone. J Mater Chem 2004, 14 (14), 2115–2123. 10.1039/B402005G.

(47) Conward, M.; Samuel, J. Machining Characteristics of the Haversian and Plexiform Components of Bovine Cortical Bone. J. Mech. Behav. Biomed. Mater. 2016, 60, 525–534. 10.1016/j.jmbbm.2016.03.017.

(48) Granke, M.; Gourrier, A.; Rupin, F.; Raum, K.; Peyrin, F.; Burghammer, M.; Saïed, A.; Laugier, P. Microfibril Orientation Dominates the Microelastic Properties of Human Bone Tissue at the Lamellar Length Scale. PLoS ONE 2013, 8 (3), e58043. 10.1371/journal.pone.0058043.

(49) Xu, J.; Rho, J. Y.; Mishra, S. R.; Fan, Z. Atomic Force Microscopy and Nanoindentation Characterization of Human Lamellar Bone Prepared by Microtome Sectioning and Mechanical Polishing Technique. J. Biomed. Mater. Res. A 2003, *67A* (3), 719–726. 10.1002/jbm.a.10109.

(50) Hengsberger, S.; Kulik, A.; Zysset, P. Nanoindentation Discriminates the Elastic Properties of Individual Human Bone Lamellae under Dry and Physiological Conditions. Bone 2002, 30 (1), 178–184. 10.1016/S8756-3282(01)00624-X.

(51) Carnelli, D.; Vena, P.; Dao, M.; Ortiz, C.; Contro, R. Orientation and Size-Dependent Mechanical Modulation within Individual Secondary Osteons in Cortical Bone Tissue. J. R. Soc. Interface 2013, 10 (81), 20120953. 10.1098/rsif.2012.0953.

(52) Gupta, H. S.; Stachewicz, U.; Wagermaier, W.; Roschger, P.; Wagner, H. D.; Fratzl, P. Mechanical Modulation at the Lamellar Level in Osteonal Bone. J. Mater. Res. 2006, 21 (8), 1913–1921. 10.1557/jmr.2006.0234.

(53) Puchegger, S.; Fix, D.; Pilz-Allen, C.; Roschger, P.; Fratzl, P.; Weinkamer, R. The Role of Angular Reflection in Assessing Elastic Properties of Bone by Scanning Acoustic Microscopy. J. Mech. Behav. Biomed. Mater. 2014, 29, 438–450. 10.1016/j.jmbbm.2013.10.004.

(54) Varga, P.; Pacureanu, A.; Langer, M.; Suhonen, H.; Hesse, B.; Grimal, Q.; Cloetens, P.; Raum, K.; Peyrin, F. Investigation of the Three-Dimensional Orientation of Mineralized Collagen Fibrils in Human Lamellar Bone Using Synchrotron X-Ray Phase Nano-Tomography. Acta Biomater. 2013, 9 (9), 8118–8127. 10.1016/j.actbio.2013.05.015.

(55) Raguin, E.; Rechav, K.; Shahar, R.; Weiner, S. Focused Ion Beam-SEM 3D Analysis of Mineralized Osteonal Bone: Lamellae and Cement Sheath Structures. Acta Biomater. 2021, 121, 497–513. 10.1016/j.actbio.2020.11.002.

(56) Gomez, S.; Rizzo, R.; Pozzi-Mucelli, M.; Bonucci, E.; Vittur, F. Zinc Mapping in Bone Tissues by Histochemistry and Synchrotron Radiation–Induced x-Ray Emission: Correlation with the Distribution of Alkaline Phosphatase. Bone 1999, 25 (1), 33–38. 10.1016/S8756-3282(99)00102-7.

(57) Pemmer, B.; Roschger, A.; Wastl, A.; Hofstaetter, J. G.; Wobrauschek, P.; Simon, R.; Thaler, H. W.; Roschger, P.; Klaushofer, K.; Streli, C. Spatial Distribution of the Trace Elements Zinc, Strontium and Lead in Human Bone Tissue. Bone 2013, 57 (1), 184–193. 10.1016/j.bone.2013.07.038.

(58) Zizak, I.; Roschger, P.; Paris, O.; Misof, B. M.; Berzlanovich, A.; Bernstorff, S.; Amenitsch, H.; Klaushofer, K.; Fratzl, P. Characteristics of Mineral Particles in the Human Bone/Cartilage Interface. J. Struct. Biol. 2003, 141 (3), 208–217. 10.1016/S1047-8477(02)00635-4.

(59) Voltolini, M.; Wenk, H.-R.; Gomez Barreiro, J.; Agarwal, S. C. Hydroxylapatite Lattice Preferred Orientation in Bone: A Study of Macaque, Human and Bovine Samples. J. Appl. Crystallogr. 2011, 44 (5), 928–934. 10.1107/S0021889811024344.

(60) Wenk, H.-R.; Grigull, S. Synchrotron Texture Analysis with Area Detectors. J. Appl. Crystallogr. 2003, 36 (4), 1040–1049. 10.1107/S0021889803010136.

(61) Schemenz, V. Correlations between Osteocyte Lacuno-Canalicular Network and Material Characteristics in Bone Adaptation and Regeneration, 2022. https://publishup.uni-potsdam.de/frontdoor/index/index/docId/55959.

(62) Wittig, N. K.; Palle, J.; Østergaard, M.; Frølich, S.; Birkbak, M. E.; Spiers, K. M.; Garrevoet, J.; Birkedal, H. Bone Biomineral Properties Vary across Human Osteonal Bone. ACS Nano 2019, 13 (11), 12949–12956. 10.1021/acsnano.9b05535.

(63) Pabisch, S.; Wagermaier, W.; Zander, T.; Li, C.; Fratzl, P. Imaging the Nanostructure of Bone and Dentin Through Small- and Wide-Angle X-Ray Scattering. In Methods in Enzymology; Elsevier, 2013; Vol. 532, pp 391–413. 10.1016/B978-0-12-416617-2.00018-7.

(64) Roschger, A.; Wagermaier, W.; Gamsjaeger, S.; Hassler, N.; Schmidt, I.; Blouin, S.; Berzlanovich, A.; Gruber, G. M.; Weinkamer, R.; Roschger, P.; Paschalis, E. P.; Klaushofer, K.; Fratzl, P. Newly Formed and Remodeled Human Bone Exhibits Differences in the Mineralization Process. Acta Biomater. 2020, 104, 221–230. 10.1016/j.actbio.2020.01.004.

(65) Hoerth, R. M.; Kerschnitzki, M.; Aido, M.; Schmidt, I.; Burghammer, M.; Duda, G. N.; Fratzl, P.; Willie, B. M.; Wagermaier, W. Correlations between Nanostructure and Micromechanical Properties of Healing Bone. J. Mech. Behav. Biomed. Mater. 2018, 77, 258–266. 10.1016/j.jmbbm.2017.08.022.

(66) Reznikov, N.; Steele, J. A. M.; Fratzl, P.; Stevens, M. M. A Materials Science Vision of Extracellular Matrix Mineralization. Nat. Rev. Mater. 2016, 1 (8), 16041. 10.1038/natrevmats.2016.41.

(67) Sims, N. A.; Martin, T. J. Coupling the Activities of Bone Formation and Resorption: A Multitude of Signals within the Basic Multicellular Unit. BoneKEy Rep. 2014, 3. 10.1038/bonekey.2013.215.

(68) Poundarik, A. A.; Boskey, A.; Gundberg, C.; Vashishth, D. Biomolecular Regulation, Composition and Nanoarchitecture of Bone Mineral. Sci. Rep. 2018, 8 (1), 1191. 10.1038/s41598-018-19253-w.

(69) Tang, T.; Landis, W.; Blouin, S.; Bertinetti, L.; Hartmann, M. A.; Berzlanovich, A.; Weinkamer, R.; Wagermaier, W.; Fratzl, P. Subcanalicular Nanochannel Volume Is Inversely Correlated With Calcium Content in Human Cortical Bone. J. Bone Miner. Res. 2022, jbmr.4753. 10.1002/jbmr.4753.

(70) Akkus, O.; Polyakova-Akkus, A.; Adar, F.; Schaffler, M. B. Aging of Microstructural Compartments in Human Compact Bone*. J. Bone Miner. Res. 2003, 18 (6), 1012–1019. 10.1359/jbmr.2003.18.6.1012.

(71) Miyaji, F.; Kono, Y.; Suyama, Y. Formation and Structure of Zinc-Substituted Calcium Hydroxyapatite. Mater. Res. Bull. 2005, 40 (2), 209–220. 10.1016/j.materresbull.2004.10.020.

(72) Paschalis, E. P.; Gamsjaeger, S.; Klaushofer, K. Vibrational Spectroscopic Techniques to Assess Bone Quality. Osteoporos. Int. 2017, 28 (8), 2275–2291. 10.1007/s00198-017-4019-y.

(73) Ping, H.; Wagermaier, W.; Horbelt, N.; Scoppola, E.; Li, C.; Werner, P.; Fu, Z.; Fratzl, P. Mineralization Generates Megapascal Contractile Stresses in Collagen Fibrils. Science 2022, 376 (6589), 188–192. 10.1126/science.abm2664.

(74) Schemenz, V.; Scoppola, E.; Zaslansky, P.; Fratzl, P. Bone Strength and Residual Compressive Stress in Apatite Crystals. J. Struct. Biol. 2024, 216 (4), 108141. 10.1016/j.jsb.2024.108141.

(75) Bar-On, B.; Wagner, H. D. New Insights into the Young’s Modulus of Staggered Biological Composites. Mater. Sci. Eng. C 2013, 33 (2), 603–607. 10.1016/j.msec.2012.10.003.

(76) Vaughan, T. J.; McCarthy, C. T.; McNamara, L. M. A Three-Scale Finite Element Investigation into the Effects of Tissue Mineralisation and Lamellar Organisation in Human Cortical and Trabecular Bone. J. Mech. Behav. Biomed. Mater. 2012, 12, 50–62. 10.1016/j.jmbbm.2012.03.003.

(77) Zhang, S.; Bach-Gansmo, F. L.; Xia, D.; Besenbacher, F.; Birkedal, H.; Dong, M. Nanostructure and Mechanical Properties of the Osteocyte Lacunar-Canalicular Network-Associated Bone Matrix Revealed by Quantitative Nanomechanical Mapping. Nano Res. 2015, 8 (10), 3250–3260. 10.1007/s12274-015-0825-8.

(78) McCutchen, C. W. Do Mineral Crystals Stiffen Bone by Straitjacketing Its Collagen? J. Theor. Biol. 1975, 51 (1), 51–58. 10.1016/0022-5193(75)90138-1.

(79) Yerramshetty, J. S.; Akkus, O. The Associations between Mineral Crystallinity and the Mechanical Properties of Human Cortical Bone. Bone 2008, 42 (3), 476–482. 10.1016/j.bone.2007.12.001.

(80) Razi, H.; Predan, J.; Fischer, F. D.; Kolednik, O.; Fratzl, P. Damage Tolerance of Lamellar Bone. Bone 2020, 130, 115102. 10.1016/j.bone.2019.115102.

(81) Li, S.; Abdel-Wahab, A.; Demirci, E.; Silberschmidt, V. V. Fracture Process in Cortical Bone: X-FEM Analysis of Microstructured Models. Int. J. Fract. 2013, 184 (1–2), 43–55. 10.1007/s10704-013-9814-7.

(82) Jonvaux, J.; Hoc, T.; Budyn, É. Analysis of Micro Fracture in Human Haversian Cortical Bone under Compression. Int. J. Numer. Methods Biomed. Eng. 2012, 28 (9), 974–998. 10.1002/cnm.2478.

(83) Wang, M.; Li, S.; Scheidt, A. vom; Qwamizadeh, M.; Busse, B.; Silberschmidt, V. V. Numerical Study of Crack Initiation and Growth in Human Cortical Bone: Effect of Micro-Morphology. Eng. Fract. Mech. 2020, 232, 107051. 10.1016/j.engfracmech.2020.107051.

(84) O’Brien, F. J.; Taylor, D.; Lee, T. C. The Effect of Bone Microstructure on the Initiation and Growth of Microcracks. J. Orthop. Res. 2005, 23 (2), 475–480. 10.1016/j.orthres.2004.08.005.

(85) Yaraghi, N. A.; Guarín-Zapata, N.; Grunenfelder, L. K.; Hintsala, E.; Bhowmick, S.; Hiller, J. M.; Betts, M.; Principe, E. L.; Jung, J.; Sheppard, L.; Wuhrer, R.; McKittrick, J.; Zavattieri, P. D.; Kisailus, D. A Sinusoidally Architected Helicoidal Biocomposite. Adv. Mater. 2016, 28 (32), 6835–6844. 10.1002/adma.201600786.

(86) Weaver, J. C.; Milliron, G. W.; Miserez, A.; Evans-Lutterodt, K.; Herrera, S.; Gallana, I.; Mershon, W. J.; Swanson, B.; Zavattieri, P.; DiMasi, E.; Kisailus, D. The Stomatopod Dactyl Club: A Formidable Damage-Tolerant Biological Hammer. Science 2012, 336 (6086), 1275–1280. 10.1126/science.1218764.

(87) Delaunois, Y.; Tits, A.; Grossman, Q.; Smeets, S.; Malherbe, C.; Eppe, G.; Van Lenthe, G. H.; Ruffoni, D.; Compère, P. Design Strategies of the Mantis Shrimp Spike: How the Crustacean Cuticle Became a Remarkable Biological Harpoon. Nat. Sci. 2023, 3 (3), e20220060. 10.1002/ntls.20220060.

(88) Stockhausen, K. E.; Qwamizadeh, M.; Wölfel, E. M.; Hemmatian, H.; Fiedler, I. A. K.; Flenner, S.; Longo, E.; Amling, M.; Greving, I.; Ritchie, R. O.; Schmidt, F. N.; Busse, B. Collagen Fiber Orientation Is Coupled with Specific Nano-Compositional Patterns in *Dark* and *Bright* Osteons Modulating Their Biomechanical Properties. ACS Nano 2021, 15 (1), 455–467. 10.1021/acsnano.0c04786.

(89) Granke, M.; Does, M. D.; Nyman, J. S. The Role of Water Compartments in the Material Properties of Cortical Bone. Calcif. Tissue Int. 2015, 97 (3), 292–307. 10.1007/s00223-015-9977-5.

(90) Bushby, A. J.; Ferguson, V. L.; Boyde, A. Nanoindentation of Bone: Comparison of Specimens Tested in Liquid and Embedded in Polymethylmethacrylate. J Mater Res 2004, 19 (1).

(91) Roschger, P.; Paschalis, E. P.; Fratzl, P.; Klaushofer, K. Bone Mineralization Density Distribution in Health and Disease. Bone 2008, 42 (3), 456–466. 10.1016/j.bone.2007.10.021.

(92) Roschger, P.; Fratzl, P.; Eschberger, J.; Klaushofer, K. Validation of Quantitative Backscattered Electron Imaging for the Measurement of Mineral Density Distribution in Human Bone Biopsies. Bone 1998, 23 (4), 319–326. 10.1016/S8756-3282(98)00112-4.

(93) Roschger, P.; Eschberger, J.; Jr, H. P. Formation of Ultracracks in Methacrylate-Embedded Undecalcified Bone Samples by Exposure to Aqueous Solutions.

(94) Lukas, C.; Kollmannsberger, P.; Ruffoni, D.; Roschger, P.; Fratzl, P.; Weinkamer, R. The Heterogeneous Mineral Content of Bone—Using Stochastic Arguments and Simulations to Overcome Experimental Limitations. J. Stat. Phys. 2011, 144 (2), 316–331. 10.1007/s10955-011-0209-8.

(95) Hartmann, M. A.; Blouin, S.; Misof, B. M.; Fratzl-Zelman, N.; Roschger, P.; Berzlanovich, A.; Gruber, G. M.; Brugger, P. C.; Zwerina, J.; Fratzl, P. Quantitative Backscattered Electron Imaging of Bone Using a Thermionic or a Field Emission Electron Source. Calcif. Tissue Int. 2021, 109 (2), 190–202. 10.1007/s00223-021-00832-5.

(96) Campbell, S. E.; Ferguson, V. L.; Hurley, D. C. Nanomechanical Mapping of the Osteochondral Interface with Contact Resonance Force Microscopy and Nanoindentation. Acta Biomater. 2012, 8 (12), 4389–4396. 10.1016/j.actbio.2012.07.042.

(97) Donnelly, E.; Baker, S. P.; Boskey, A. L. Effects of Surface Roughness and Maximum Load on the Mechanical Properties of Cancellous Bone Measured by Nanoindentation. 2006.

(98) Bobji, M. S.; Biswas, S. K. Estimation of Hardness by Nanoindentation of Rough Surfaces. J. Mater. Res. 1998, 13 (11), 3227–3233. 10.1557/JMR.1998.0438.

(99) Oliver, W. C.; Pharr, G. M. An Improved Technique for Determining Hardness and Elastic Modulus Using Load and Displacement Sensing Indentation Experiments. J. Mater. Res. 1992, 7 (6), 1564–1583. 10.1557/JMR.1992.1564.

(100) Ascenzi, M.-G.; Ascenzi, A.; Benvenuti, A.; Burghammer, M.; Panzavolta, S.; Bigi, A. Structural Differences between “Dark” and “Bright” Isolated Human Osteonic Lamellae. J. Struct. Biol. 2003, 141 (1), 22–33. 10.1016/S1047-8477(02)00578-6.

(101) Benecke, G.; Wagermaier, W.; Li, C.; Schwartzkopf, M.; Flucke, G.; Hoerth, R.; Zizak, I.; Burghammer, M.; Metwalli, E.; Müller-Buschbaum, P.; Trebbin, M.; Förster, S.; Paris, O.; Roth, S. V.; Fratzl, P. A Customizable Software for Fast Reduction and Analysis of Large X-Ray Scattering Data Sets: Applications of the New *DPDAK* Package to Small-Angle X-Ray Scattering and Grazing-Incidence Small-Angle X-Ray Scattering. J. Appl. Crystallogr. 2014, 47 (5), 1797–1803. 10.1107/S1600576714019773.

(102) Fratzl, P.; Schreiber, S.; Klaushofer, K. Bone Mineralization as Studied by Small-Angle X-Ray Scattering. Connect. Tissue Res. 1996, 34 (4), 247–254. 10.3109/03008209609005268.

(103) Fratzl, P. Statistical Model of the Habit and Arrangement of Mineral Crystals in the Collagen of Bone. J. Stat. Phys. 1994, 77 (1–2), 125–143. 10.1007/BF02186835.

(104) Danilchenko, S. N.; Kukharenko, O. G.; Moseke, C.; Protsenko, I. Yu.; Sukhodub, L. F.; Sulkio-Cleff, B. Determination of the Bone Mineral Crystallite Size and Lattice Strain from Diffraction Line Broadening. Cryst. Res. Technol. 2002, 37 (11), 1234–1240. 10.1002/1521-4079(200211)37:11<1234::AID-CRAT1234>3.0.CO;2-X.

(105) Handschin, R. G.; Stern, W. B. X-Ray Diffraction Studies on the Lattice Perfection of Human Bone Apatite (Crista Iliaca). 16 (4).

(106) Rinnerthaler, S.; Roschger, P.; Jakob, H. F.; Nader, A.; Klaushofer, K.; Fratzl, P. Scanning Small Angle X-Ray Scattering Analysis of Human Bone Sections. Calcif. Tissue Int. 1999, 64 (5), 422–429. 10.1007/PL00005824.

